# FAIRsharing, a cohesive community approach to the growth in standards, repositories and policies

**DOI:** 10.1101/245183

**Authors:** Susanna-Assunta Sansone, Peter McQuilton, Philippe Rocca-Serra, Alejandra Gonzalez-Beltran, Massimiliano Izzo, Allyson L. Lister, Milo Thurston, And the FAIRsharing community (https://fairsharing.org/communities):

**Author notes:** Disclaimer: The views presented in this article do not necessarily reflect current or future opinion or policy of the US Food and Drug Administration. Any mention of commercial products is for clarification and not intended as an endorsement.

## Abstract

In this modern, data-driven age, governments, funders and publishers expect greater transparency and reuse of research data, as well as greater access to and preservation of the data that supports research findings. Community-developed standards, such as those for the identification^1^ and reporting^2^ of data, underpin reproducible and reusable research, aid scholarly publishing, and drive both the discovery and evolution of scientific practice. The number of these standardization efforts, driven by large organizations or at the grass root level, has been on the rise since the early 2000s. Thousands of community-developed standards are available (across all disciplines), many of which have been created and/or implemented by several thousand data repositories. Nevertheless, their uptake by the research community, however, has been slow and uneven. This is mainly because investigators lack incentives to follow and adopt standards. The situation is exacerbated if standards are not promptly implemented by databases, repositories and other research tools, or endorsed by infrastructures. Furthermore, the fragmentation of community efforts results in the development of arbitrarily different, incompatible standards. In turn, this leads to standards becoming rapidly obsolete in fast-evolving research areas.

As with any other digital object, standards, databases and repositories are dynamic in nature, with a ‘life cycle’ that encompasses formulation, development and maintenance; their status in this cycle may vary depending on the level of activity of the developing group or community. There is an urgent need for a service that enhances the information available on the evolving constellation of heterogeneous standards, databases and repositories, guides users in the selection of these resources, and that works with developers and maintainers of these resources to foster collaboration and promote harmonization. Such an informative and educational service is vital to reduce the knowledge gap among those involved in producing, managing, serving, curating, preserving, publishing or regulating data. A diverse set of stakeholders-representing academia, industry, funding agencies, standards organizations, infrastructure providers and scholarly publishers— both national and domain-specific as well global and general organizations— have come together as a community, representing the core adopters, advisory board members, and/or key collaborators of the FAIRsharing resource. Here, we introduce its mission and community network. We present an evaluation of the standards landscape, focusing on those for reporting data and metadata - the most diverse and numerous of the standards - and their implementation by databases and repositories. We report on the ongoing challenge to recommend resources, and we discuss the importance of making standards invisible to the end users. We report on the ongoing challenge to recommend resources, and we discuss the importance of making standards invisible to the end users. We present guidelines that highlight the role each stakeholder group must play to maximize the visibility and adoption of standards, databases and repositories.

## Mapping the landscape and tracking evolution

Working with and for data producers and consumers, and leveraging on our large network of international collaborators we have iteratively^2^,^3^,^4^ developed FAIRsharing (https://fairsharing.org), an informative and educational resource that describes and interlinks community-driven standards, databases, repositories, and data policies. As of 10th of July 2018, FAIRsharing has over 2352 records: 1173 standards, 1071 data repositories, 112 data policies (of which 80 are from journals and publishers and 22 from funders). These records focus mainly on the life, agricultural, environmental, biomedical and health sciences, but FAIRsharing is progressively expanding to cover other disciplines. However, quantity is not the end goal. The richness and accuracy of each record are our priorities.

Using community participation, the FAIRsharing team precisely curates information on standards employed for the identification, citation and reporting of data and metadata, via four standards subtypes. Minimum reporting guidelines, also known as guiding principles or checklists, outline the necessary and sufficient information vital for contextualizing and understanding a digital object. Terminology artifacts or *semantics*, ranging from dictionaries to ontologies, provide definitions and unambiguous identification for concepts and objects. Models and formats define the structure and relationship of information for a conceptual model or schema, and include transmission formats to facilitate the exchange of data between different systems. Identifier schema are formal systems for resources and other digital objects that allow their unique and unambiguous identification. These standards range from generic and multi-disciplinary, to standards that are tailored for specific disciplines. FAIRsharing monitors their evolution, implementation in databases and repositories, and recommendation by journal and funder data policies.

Producers of standards, databases and repositories are able to claim the record(s) for the resource(s) they maintain or have developed; this functionality allows them to gain personal recognition and ensures that the description is accurate and up-to-date. All records and related updates by the maintainers are checked by a FAIRsharing curator. Conversely if a record is updated by a FAIRsharing curator, an email notification is sent to the record claimant which minimizes the introduction of inaccuracies. In communication with the community behind each resource, FAIRsharing assigns indicators to show the status in the resource’s life cycle: ‘Ready’ for use, ‘In Development’, ‘Uncertain’ (when any attempt to reach out to the developing community has failed), and ‘Deprecated’ (when the community no longer mandates its use, together with an explanation where available).

To make standards, databases, repositories and data policies more discoverable and citable, we mint digital object identifiers (DOIs) for each record, which provides a persistent and unique identifier to enable referencing of these resources. In addition, the maintainers of each record can be linked with their Open Research and Contributor IDentifier (ORCID) profile (https://orcid.org). Citing a FAIRsharing record for a standard, database and repository offers a unique, at-a-glance view of all descriptors and indicators pertaining to a resource, as well as any evidence of adoption or endorsement by a data policy or organisation. Referencing the record together with the resource’s main paper (which provides a snapshot of its status at a given time) provides a complete reference for a resource. FAIRsharing has its own record to serve this very purpose: https://doi.org/10.25504/FAIRsharing.2abjs5.

Working with and for the community, FAIRsharing collects the necessary information to ensure that standards, databases, repositories and data policies are aligned with the FAIR data principles^5^. It ensures these resources are Findable (e.g., by providing persistent and unique identifiers, functionalities to register, claim, maintain, inter-link, search and discover them), Accessible (e.g., identifying their level of openness and/or licence type), encouraged to be Interoperable (e.g., highlighting which repositories implement the same standards to structure and exchange data), and Reusable (e.g., knowing the coverage of a standard and its level of endorsement by a number of repositories should encourage its use or extension in neighbouring domains, rather than reinvention). With the goal of being an interoperable component in the ecosystem of other services, FAIRsharing collaborates with many other infrastructure resources to cross-link each record to other registries, as well as within major FAIR-driven global initiatives, research and infrastructure programmes, many of which are generic and cross-disciplinary. Exemplars are listed in **Box1**, along with the roles that FAIRsharing plays. A ‘live’, updated list is maintained at https://fairsharing.org/communities/activities. An example is the FAIR Metrics working group (http://fairmetrics.org)^6^, where we work to guide producers of standards, databases and repositories to assess the level of FAIRness of their resource. We will develop measurable indicators, which will be progressively implemented in the FAIRsharing registry. The content within FAIRsharing is licensed via the Creative Commons Attribution ShareAlike license 4.0 (CC BY-SA 4.0); the SA
clause enhances the open heritage and aims to create a larger open commons, ensuring the downstream users share back

**Box 1:** Exemplar research infrastructure programmes and umbrella organisations that FAIRsharing is part of and working with.

- **CODATA** (http://www.codata.org): the interdisciplinary scientific Committee on Data of the International Science Council, CODATA promotes global collaboration to improve the availability and usability of data for all areas of research. FAIRsharing is an active contributor to the activities of the CODATA Data Integration and Interoperability Initiative (http://dataintegration.codata.org).
- **ELIXIR** (https://www.elixir-europe.org): an intergovernmental organisation that brings together life science resources from across Europe and beyond to build a sustainable infrastructure to support life science research and its translation to medicine and the environment, the bio-industries and society. FAIRsharing is a service of its Interoperability Platform.
- **EOSC Pilot** (https://eoscpilot.eu): the European Open Science Cloud (EOSC) Pilot that supports the exploration of a virtual environment with open and seamless services for storage, management, analysis and re-use of research data, across borders and scientific disciplines by federating existing scientific data infrastructures. FAIRsharing is a core element of its proposed metadata catalogues strategy.
- **FORCE11** (https://www.force11.org): a community of scholars, librarians, archivists, publishers and research funders that aims to bring about a change in modern scholarly communications through the effective use of information technology. FAIRsharing has a working group in FORCE11 and works with its Data Citation Implementation Pilot, including journals and publishers, to identify criteria and develop tools for the selection of databases and repositories.
- **GO-FAIR** (http://go-fair.org): the Global and Open (GO) FAIR initiative for the practical implementation of the EOSC vision as part of a global Internet of FAIR Data & Services. FAIRsharing is part of several bottom up community efforts (called Implementation Networks) to build an ecosystem of FAIR services (http://fairsharing.fairdata.solutions).
- **ISO** (https://www.iso.org): the International Organization for Standardization (ISO) is a non-governmental international organization, with a membership of 161 national standards bodies, developing standards via Technical Committees (TCs). The TC 276 focuses on the biotechnology process and its Working Group 5 is in charge of the standards for Data Processing and Integration. FAIRsharing works to provide a trackable list of the standards endorsed by the TC 276 Biotechnology WG5.
- **NIH Data Commons** (https://commonfund.nih.gov/commons): the US National Institutes of Health (NIH; Bethesda, MD) Data Commons Pilot Phase explores the design and creation of a shared virtual space where scientists can work with the digital objects of biomedical research such as data and analytical tools. FAIRsharing is functional element of a distributed FAIRness assessment tool kit, including the FAIRshake system to evaluate digital objects.
- **RDA** (https://www.rd-alliance.org): an international organization that focuses on the development of infrastructure and community activities to reduce barriers to data sharing and promote the acceleration of data driven innovation worldwide. FAIRsharing has a working group in RDA and works with other RDA activities, e.g., to define and implement a common framework for journal and publisher research data journal policy; and to connect FAIRsharing to data management plans tools.

## We say we need standards, but do we use them?

The scientific community, funders and publishers all endorse the concept that common data and metadata standards underpin data reproducibility, ensuring that the relevant elements of a dataset are reported and shared consistently and meaningfully. However, navigating through the many standards available can be discouraging and often unappealing for prospective users. Bound within a particular discipline or domain, reporting standards are fragmented, with gaps and duplications, thereby limiting their combined used. Although standards should stand alone, they should also function well together, especially to better support multi-dimensional data but also the aggregation of pre-existing datasets from one or more disciplines or domains. Although standards should stand alone, they should also function well together, especially to better support multi-dimensional data but also the aggregation of pre-existing datasets from one or more disciplines or domains. Understanding how they work or how to comply with them takes time and effort. Measuring the uptake of standards, however, is not trivial, and achieving a full picture is practically impossible.

FAIRsharing provides a snapshot of the standards landscape, which is dynamic and will continue to evolve as we engage with more communities and verify the information we house, add new resources, track their life-cycle status and usage in databases and repositories, and link out to examples of training material. FAIRsharing also plays a fundamental role in the activation of the decision-making chain, which is is an essential step towards fostering the wider adoption of standards. When a standard is mature and appropriate standard-compliant systems become available, such as databases and repositories, these must then be channelled to the relevant stakeholder community, who in turn must recommend them (e.g., in data policies) or use them (e.g. to define a data management plan) to facilitate a high-quality research cycle.

As of 10th of July 2018, there are a total of 1173 community standards, 793 of which are specific to the life, agricultural, environmental, biomedical and health sciences, and 45 are generic and multi-disciplinary. 124 reporting guidelines (out of 144), 653 terminology artifacts (out of 737), and 6 identifier schemas (out of 7) are mature and tagged as ‘Ready’ for use. **Table 1** displays the 12 data and metadata standard records, corresponding to the top ten positions, that were most accessed on FAIRsharing during 2017. This ranking, however, shows no direct correlation with the level of standard adoption (by journal and funder data policies, or by databases and repositories) and is perhaps due to their popularity within their direct domain. The ranking is also very variable and can change substantially from year to year, most likely reflecting the activity of their respective research communities and when a standard is actively in development.

**Table 1.**
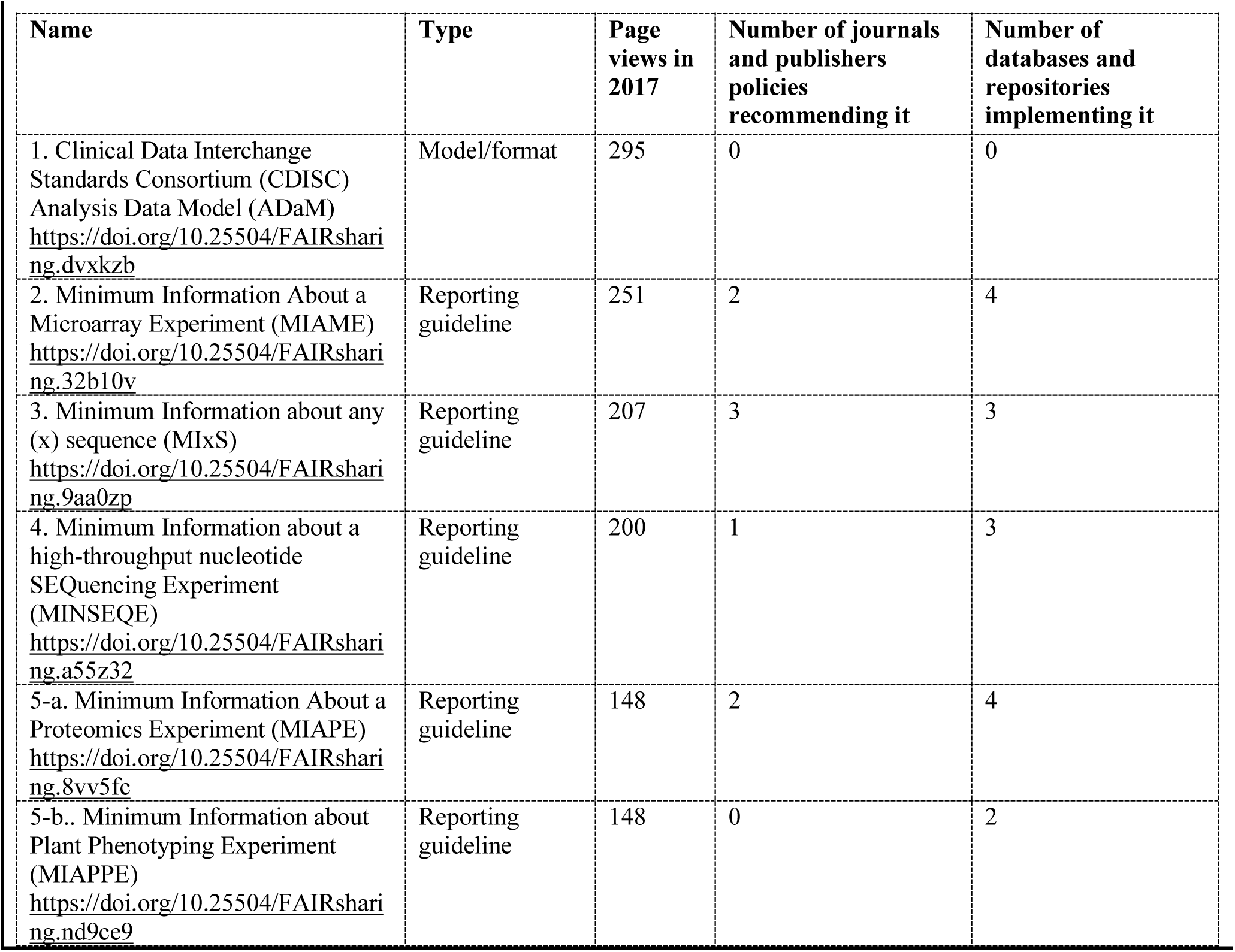
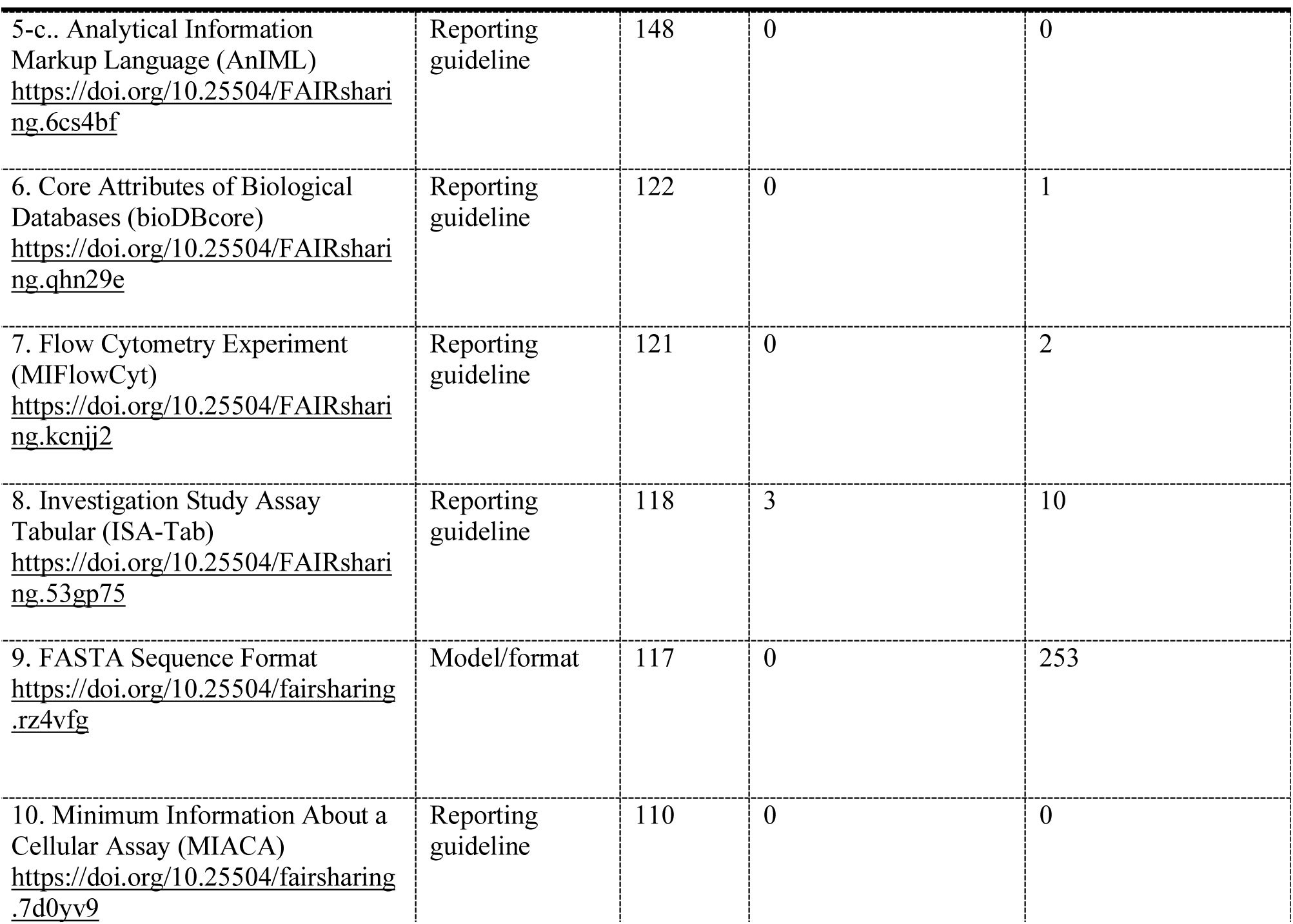
The 12 data and metadata standards in the top ten positions (all tagged as “Ready”) are ranked according to the page views in 2017; in addition the number of data policies that recommend them, along with the number of databases and repositories that implement them are given.

**Table 2** displays the top ten data and metadata standard records that have been implemented by databases and repositories, providing a realistic measure of the use of data and metadata standards to annotate, structure and share datasets. Surprisingly, with the exception of one (the NCBI Taxonomy, a terminology artifact for taxonomic information: https://doi.org/10.25504/FAIRsharing.fi07xj). none of the other nine standards are explicitly recommended in journals and databases’ data policies, including the standard most implemented by databases and repositories (the FASTA Sequence Format, model/format for representing either nucleotide sequences or peptide sequences: https://doi.org/10.25504/FAIRsharing.rz4vfg). This omission can probably be explained by the fact that, created in 1985, this is a *de facto* standard that every sequence database and repositories implements by default, thus becoming (positively) ‘invisible’ to users, including publishers/journals.

**Table 2.** As of the 10th of July 2018, the top ten data and metadata standards (all of which are tagged as
“Ready”) ranked according to the number of implementations by databases and repositories; in addition the number of data policies that recommend them, along with the number of page views in 2017 are provided.

| Name | Type | Number of databases and repositories implementing it | Number of journals and publishers policies recommending it | Page views in 2017 |
| --- | --- | --- | --- | --- |
| 1. FASTA Sequence Format<br>https://doi.org/10.25504/FAIRsharing.rz4vfg | Model/format | 253 | 0 | 117 |
| 2. Gene Ontology (GO)<br>https://doi.org/10.25504/FAIRsharing.6xq0ee | Terminology artifact | 144 | 0 | 69 |
| 3. Protein Data Bank (PDB) Format<br>https://doi.org/10.25504/FAIRsharing.9y4cqW | Model/format | 58 | 0 | 25 |
| 4. Generic Feature Format Version 3 (GFF3)<br>https://doi.org/10.25504/FAIRsharing.dnk0f6 | Model/format | 47 | 0 | 12 |
| 5. Chemical Entities of Biological Interest (ChEBI)<br>https://doi.org/10.25504/FAIRsharing.62qk8w | Terminology artifact | 30 | 0 | 49 |
| 6. Schema.org<br>https://doi.org/10.25504/FAIRsharing.hzdqz8 | Model/format | 29 | 0 | 19 |
| 7. GenBank Sequence Format<br>https://doi.org/10.25504/FAIRsharing.rg2vmt | Model/format | 28 | 0 | 19 |
| 8. NCBI Taxonomy (NCBITAXON)<br>https://doi.org/10.25504/FAIRsharing.fj07xj | Terminology artifact | 25 | 3 | 91 |
| 9. Sequence Ontology (SO)<br>https://doi.org/10.25504/FAIRsharing.6bc7h9 | Terminology artifact | 22 | 0 | 25 |
| 10. Molecular Interaction Tabular (MITAB)<br>https://doi.org/10.25504/FAIRsharing.ve0710 | Model/format | 19 | 0 | 26 |

To understand how journals and publishers select which resource to recommend, we have worked closely with the editors of the following journals/publishers, whose total of 13 data policies (as of July 10th 2018) are quite well developed (‘live’ list is at: https://fairsharing.org/recommendations): EMBO Press, *F1000Research* (including five F1000-powered publication platforms - such as the funder-related Bill & Melinda Gates Foundation’s *Gates Open Research* and Wellcome Trust’s *Wellcome Open Research*), Oxford University Press’ *GigaScience*, PLOS, Elsevier and Springer Nature’s BioMed Central and *Scientific Data* (which extends to other Scientific- and Nature-titled journals). As of 10th of July 2018, the data policies of these journals/publishers recommend a total of 33 standards: 18 reporting guidelines, 8 terminology artifacts and 7 models/formats, as shown in **Table 3** (a ‘live’ list can be viewed at: https://fairsharing.org/article/live_list_standards_in_policies). Surprisingly, out of these 33 recommended standards, only one (the NCBI Taxonomy) is in the top ten standards most implemented by databases and repositories (as shown in Table 1), whilst one third (ten reporting guidelines and one terminology artifact) are not implemented.

**Table 3.** As of the 10th of July 2018, the 33 data and metadata standards in the top five positions (all of which are tagged as “Ready”) ranked according to the number of recommendations by the 13 main journals/publishers data policies; in addition, the number of databases and repositories that implement them, along with the page views in 2017, are given.

| Name | Type | Number of journals and publishers policies recommending it | Number of databases and repositories implementing it | Page views in 2017 |
| --- | --- | --- | --- | --- |
| 1-a. FORCE11 Data Citation Principles<br>https://doi.org/10.25504/FAIRsharing.9hynwc | Reporting guideline | 9 | 2 | 26 |
| 1-b. Animals in Research: Reporting In Vivo Experiments (ARRIVE)<br>https://doi.org/10.25504/FAIRsharing.t58zhj | Reporting guideline | 9 | 0 | 76 |
| 1-c. CONSOLidated standards of Reporting Trials (CONSORT)<br>https://doi.org/10.25504/FAIRsharing.gr06tm | Reporting guideline | 9 | 0 | 35 |
| 1-d. Preferred Reporting Items for Systematic reviews and Meta-Analyses (PRISMA)<br>https://doi.org/10.25504/FAIRsharing.gp3r4n | Reporting guideline | 9 | 0 | 32 |
| 1-e. Case Reports<br>https://doi.org/10.25504/FAIRsharing.zgqy0v | Reporting guideline | 9 | 0 | 18 |
| 2. DataCite Metadata Schema<br>https://doi.org/10.25504/FAIRsharing.me4qwe | Model/format | 7 | 6 | 70 |
| 3-a. NCBI Taxonomy (NCBITAXON)<br>https://doi.org/10.25504/FAIRsharing.fj07xj | Terminology artifact | 3 | 25 | 91 |
| 3-b. Investigation Study Assay Tabular (ISA-Tab)<br>https://doi.org/10.25504/FAIRsharing.53gp75 | Model/format | 3 | 9 | 124 |
| 3-c. Minimum Information about any (x) sequence (MIxS)<br>https://doi.org/10.25504/FAIRsharing.9aa0zp | Reporting guideline | 3 | 3 | 210 |
| 4-a. Minimum Information About a Microarray Experiment (MIAME)<br>https://doi.org/10.25504/FAIRsharing.32b10v | Reporting guideline | 2 | 4 | 254 |
| 4-b. Minimum Information About a Proteomics Experiment (MIAPE)<br>https://doi.org/10.25504/FAIRsharing.8vv5fc | Reporting guideline | 2 | 4 | 150 |
| 4-c. Minimum Information about a Molecular Interaction Experiment (MIMIx)<br>https://doi.org/10.25504/FAIRsharing.8z3xzh | Reporting guideline | 2 | 4 | 51 |
| 4-d. MIAME Notation in Markup Language (MINiML)<br>https://doi.org/10.25504/FAIRsharing.gaegy8 | Model/format | 2 | 3 | 43 |
| 4-e. Consolidated criteria for reporting qualitative research (COREQ)<br>https://doi.org/10.25504/FAIRsharing.6mhzhj | Reporting guideline | 2 | 0 | 20 |
| 4-f. STrengthening the Reporting of OBservational studies in Epidemiology (STROBE)<br>https://doi.org/10.25504/FAIRsharing.1mk4v9 | Reporting guideline | 2 | 0 | 20 |
| 4-g. STAndards for the Reporting of Diagnostic (STARD) accuracy<br>https://doi.org/10.25504/FAIRsharing.956df7 | Reporting guideline | 2 | 0 | 17 |
| 4-h. Consolidated Health Economic Evaluation Reporting Standards (CHEERS)<br>https://doi.org/10.25504/FAIRsharing.neny94 | Reporting guideline | 2 | 0 | 8 |
| 4-i. CONSOLIDated Standards of Reporting Trials - Official Extensions (CONSORT-OE)<br>https://doi.org/10.25504/FAIRsharing.wstthd | Reporting guideline | 2 | 0 | 6 |
| 4-l. CONSolidated Standards of Reporting Trials - Unofficial Extensions (CONSORT-UE)<br>https://doi.org/10.25504/FAIRsharing.2kq1fs | Reporting guideline | 2 | 0 | 2 |
| 5-a. Systems Biology Markup Language (SBML)<br>https://doi.org/10.25504/FAIRsharing.9qv71f | Model/format | 1 | 15 | 59 |
| 5-b. Ontology for Biomedical Investigations (OBI)<br>https://doi.org/10.25504/FAIRsharing.284e1z | Terminology artifact | 1 | 9 | 44 |
| 5-c. PSI Molecular Interaction Controlled Vocabulary (PSI-MI CV)<br>https://doi.org/10.25504/FAIRsharing.8qzmtr | Terminology artifact | 1 | 8 | 34 |
| 5-d. Experimental Factor Ontology (EFO)<br>https://doi.org/10.25504/FAIRsharing.1gr4tz | Terminology artifact | 1 | 7 | 41 |
| 5-e. mz Markup Language (mzML)<br>https://doi.org/10.25504/FAIRsharing.26dmba | Model/format | 1 | 6 | 24 |
| 5-f. Minimal Information Required In the Annotation of Models (MIRIAM)<br>https://doi.org/10.25504/FAIRsharing.ap169a | Reporting guideline | 1 | 5 | 48 |
| 5-g. Environment Ontology (EnvO)<br>https://doi.org/10.25504/FAIRsharing.azqskx | Terminology artifact | 1 | 4 | 43 |
| 5-h. Minimal Information about a high throughput SEQuencing Experiment (MINSEQE)<br>https://doi.org/10.25504/FAIRsharing.a55z32 | Reporting guideline | 1 | 3 | 200 |
| 5-i. CellML<br>https://doi.org/10.25504/FAIRsharing.50n9hc | Model/format | 1 | 3 | 41 |
| 5-l. BioAssay Ontology (BAO)<br>https://doi.org/10.25504/FAIRsharing.mye76w | Terminology artifact | 1 | 1 | 35 |
| 5-m. eagle-i Research Resource Ontology (ERO)<br>https://doi.org/10.25504/FAIRsharing.nwgyn | Terminology artifact | 1 | 1 | 15 |
| 5-n. ThermoML<br>https://doi.org/10.25504/FAIRsharing.7b0fc3 | Model/format | 1 | 1 | 10 |
| 5-o. Units Ontology (UO)<br>https://doi.org/10.25504/FAIRsharing.mjnypw | Terminology artifact | 1 | 0 | 10 |

Furthermore, these data policies recommend 184 (generalistic and domain-specific) databases and repositories. The 26 that occupy the top five positions are shown in **Table 4** (a ‘live’ list is at: https://fairsharing.org/article/live_list_databases_in_policies). As expected, this top tier includes public databases and repositories from major research and infrastructure providers from the
USA and Europe (the domain-specific UniProt Knowledgebase, for functional information on proteins, is at the top of the list with the higher number of standards implemented: https://doi.org/10.25504/FAIRsharing.s1ne3g). However, this analysis also indicates that an additional 185 standards, which are implemented by the recommended databases and repositories, are not explicitly mentioned in the journals/publishers data policies.

**Table 4.**
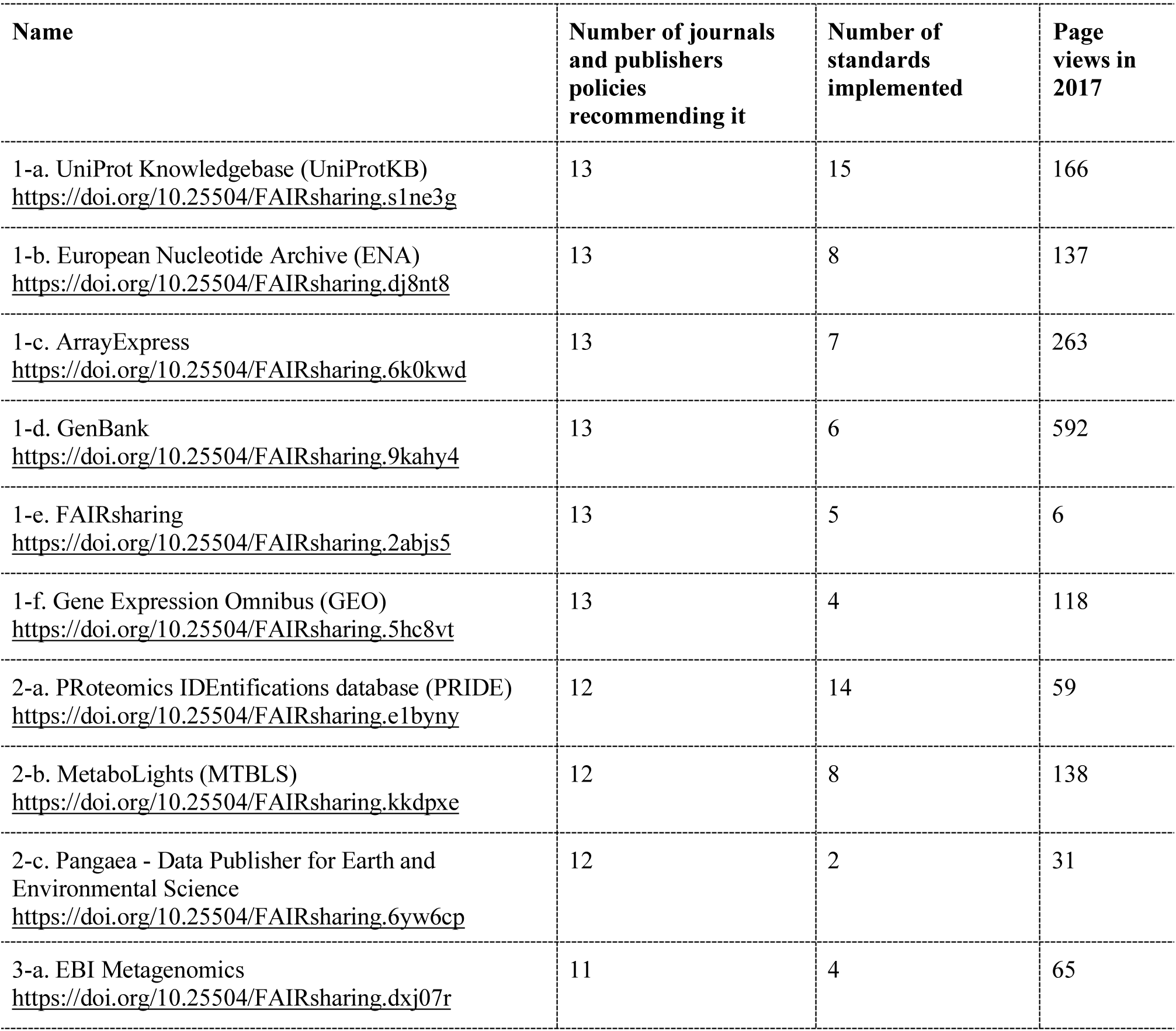
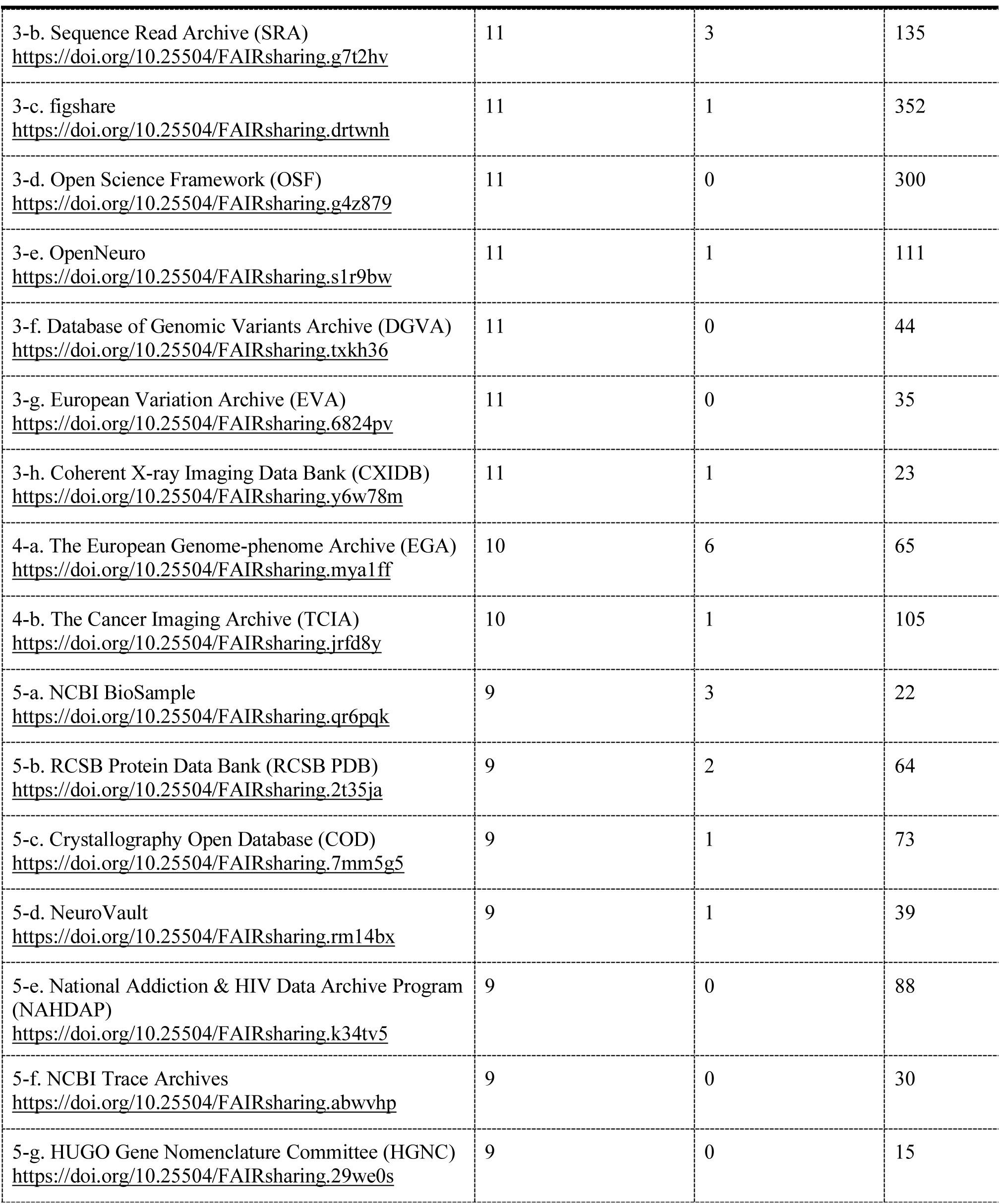
As of the 10th of July 2018, the 26 databases and repositories in the top five positions (all tagged “Ready”) ranked according to the number of recommendations by the 13 journals/publishers data policies; in addition, the number of standards these databases and repositories implement, along with the page views, is given. Due to the adoption by journals and publishers and other stakeholders, a FAIRsharing record was created at the end of 2017 (hence the low page views) to enable formal citation of the resource.

The same discrepancy between the number and type of standards that are explicitly recommended in the journals/publishers data policies, and those that are implemented by databases and repositories, is found analyzing all 80 journals and publishers data policies curated in FAIRsharing, as of 10th of July 2018 (the ‘live’ list is at: https://fairsharing.org/article/live_list_iournal_policies). Only 66 data policies mention one or more specific standards: the minimal reporting guidelines are recommended 7 times more than terminology artifacts and 4.7 times more than models/formats, even if the latter is heavily implemented by data repositories. Databases are recommended 694 times, with 181 databases recommended in total, 43 times more than models/formats.

Based on ongoing discussions with the eight journals and publishers mentioned above, along with other interested parties such as *eLife*, Taylor & Francis Group and Wiley, we understand this discrepancy in recommendation to be the consequence of a cautious approach to choosing which standard to recommend where thousands of (often competing) standards are available. It is understandable if journals/publishers do not overreach. Recommendation of a standard is often driven by the editor’s familiarity with one or more standards, notably for journals/publishers focussing on specific disciplines and areas of study, or the engagement with learned societies and researchers actively supporting and using certain standards. Generally, the community journals/publishers serve is often not familiar with standards, with many standards perceived as a hindrance to data reporting rather than help. Therefore, the current trend is for journals/publishers to recommend generalists and a core set of discipline-specific repositories - although a bigger number of (public and global, project-driven, and institution-based) databases and repositories exist, and the list of those recommended by the various organizations are very different^7^ - while very few standards, such as those for data citation standards and the minimum reporting guidelines (the metadata standards more relevant to publication), are recommended. The general opinion is that terminology artifacts and models/formats instead should emerge from a close collaboration between their developing community and the implementing repositories, and remain implicitly suggested.

FAIRsharing, therefore, plays the unique role of highlighting to journals/publishers, as well as researchers and other stakeholders, which terminology artifacts, models/formats, along other standards, each database and repository implements. This, along with community indicators of use and maturity, and emerging global certifications, is essential to inform the selection or recommendation of relevant databases and repositories. FAIRsharing aims to increase the visibility, citation and credit of these community-driven standards, databases and repositories efforts.

## The best standards are invisible and transparent

Standards for reporting of data and metadata are essential for data reuse, which drives sciences and discoveries. Minimal reporting guidelines are intended for human consumption and are usually narrative in form and therefore prone to ambiguities, making compliance and validation difficult and approximate. Many of these guidelines, however, already come with (or lead to the development of) associated models/formats and terminology artifacts, which are created to be machine readable (rather than for human consumption). These two types of standards ensure the datasets are harmonized in regard to structure, formatting and annotation, setting the foundation for the development of tools and repositories that enable transparent interpretation, verification, exchange, integrative analysis and comparison of (heterogeneous) data. The goal is to ensure the implementation of these standards in data annotation tools and data repositories, making these standards invisible to the end users.

Models/formats and terminology artifacts are essential to the implementation of the FAIR principles that put a specific emphasis on enhancing the ability of machines to automatically discover and use data. In particular, the ‘computability’ of standards is core to the development of FAIR metrics to measure the level of compliance of a given dataset against the relevant metadata descriptors. These machine-readable standards provide the necessary quantitative and verifiable measures of the degree by which data meets these reporting guidelines. The latter, on their own, would just be statements of unverifiable good intentions of compliance to given standards.

Delivering tools and practices to create standards-based templates for describing datasets smarter and faster is essential, if we are to use these standards in the authoring of metadata for the variety of data types in the life sciences and other disciplines. Community discussions are ongoing around the need for common frameworks for disciplinary research data management protocols^8^. Furthermore, research activities to deliver machine-readable standards are already being undertaken by the FAIRsharing team and collaborators^9^; all outputs will be freely shared for others to develop tools that would make it easy to check the compliance of data to standards.

## Committed to community service

The FAIRsharing mission is to increase guidance to consumers of standards, databases, repositories, and data policies, to accelerate the discovery, selection and use of these resources; and producer satisfaction in terms of resource visibility, reuse, adoption and citation. **Box 2** illustrates community-provided exemplar use cases that drive our work. This is a major undertaking, but it is a journey we are not doing alone.

**Box 2.** Exemplar on how FAIRsharing can help different stakeholders.

- Carla (**researcher**) searches FAIRsharing to identify an established repository, recognized by the journal she plans to submit to, with restricted data access to deposit her sensitive datasets, as recommended by her funder’s data policy.
- Andrea (**biocurator**) searches FAIRsharing for suitable standards to describe a set of experiments; he filters the results by disciplines, focussing on standards implemented by one or more data repositories, with available annotations tools; he also looks for examples of the most up-to-date version of the standards, and the details of a person or support group to contact.
- Alex (**standards developer**) creates and maintains a personalized collection page on FAIRsharing to list and showcase the set of standards developed by the grass-roots standard organization she is the representative of; she registers the standards and/or claims existing records added by the FAIRsharing team, vetting the descriptions and/or enhancing them by adding indicators of maturity for the standards, and indicating the repositories and tools implementing them; her grass-roots organization uses the collection to maximise the visibility of their standards, promoting adoption outside their immediate community, also favouring reuse in and extensions to other areas.
- Sam (**repository manager**) registers his data resource at FAIRsharing manually or programmatically, describing terms of deposition and access, adding information on the resource’s relationship to other repositories and use of standards, and assessing the level FAIRness of his data repository; he links the record to funding source(s) supporting the resources and the institute(s) hosting it, as well as his ORCID profile to get credit for his role as maintainer of a resource; he receives alerts if a publisher recommends his repository in a data policy; and he uses the DOI assigned to his repository record to cite the evidence of these adoption.
- Andy (**policy maker**) registers her journal’s data policy in FAIRsharing, creating and maintaining an interrelated list of the repositories and standards she recommends to the authors, to deposit and annotated data and other digital assets; she keeps her data policy up-to-date using visualization and comparison functionalities, and consulting the knowledge graph that offers a interactive view of the repositories, tools and standards, as well as receiving customized alerts, e.g. when a repository has changed its data access terms, or when a standards has been superseded by another.
- Bob (**data manager**) consults FAIRsharing when creating a data management plan to identify the most appropriate reporting guidelines, formats and terminologies for his data types, and formally cites these community standards using their DOIs and/or the ‘how to cite this record’ statements provided for each resource.
- Kyle (**librarian**) and her colleagues involved in supporting research data use FAIRsharing to: enrich educational and training material to support scholars to utilize data standards, and conform to journal and funder policies, and to develop guidance that increases capability and skills, and empowers researchers to organize and make their data FAIR.

Collaborative work is happening on many fronts. We are categorizing the records according to discipline and domain via two open application ontologies. This should facilitate more accurate browsing, discovery and selection. To improve our policy registry we are disambiguating between individual journal policies and those by publishers that encompass more journals. This will increase the number of journals covered and more accurately represent the different data policy models currently being pursued by publishers. Selection and decision making is being improved by the enrichment of indicators based on community-endorsed and discipline-specific criteria, such as FAIR metrics and FAIRness level. To maximize the ‘look-up service’ functionality, and to connect the content to other registries and tools, customizable interfaces for human as well as programmatic access to the data are being created. We are also expanding the existing network graph and creating new visually-accessible statistics (https://fairsharing.org/summary-statistics). Finally, on a monthly-basis we are highlighting featured exemplar resources, as well as adding to the informative and educational material available on FAIRsharing.

## Guidance to stakeholders

To foster a culture change within the research community into one where the use of standards, databases and repositories for FAIRer data is pervasive, we need to better promote the existence and value of these resources. First and foremost, we need to paint an accurate picture of the status quo. Several stakeholders can play catalytic roles (**Figure 1**); specifically, developers/curators of these resources; journal editors and publishers; research data management support staff, trainers and educators; societies, unions and communities alliances; funders; and finally researchers.

**Figure 1.**
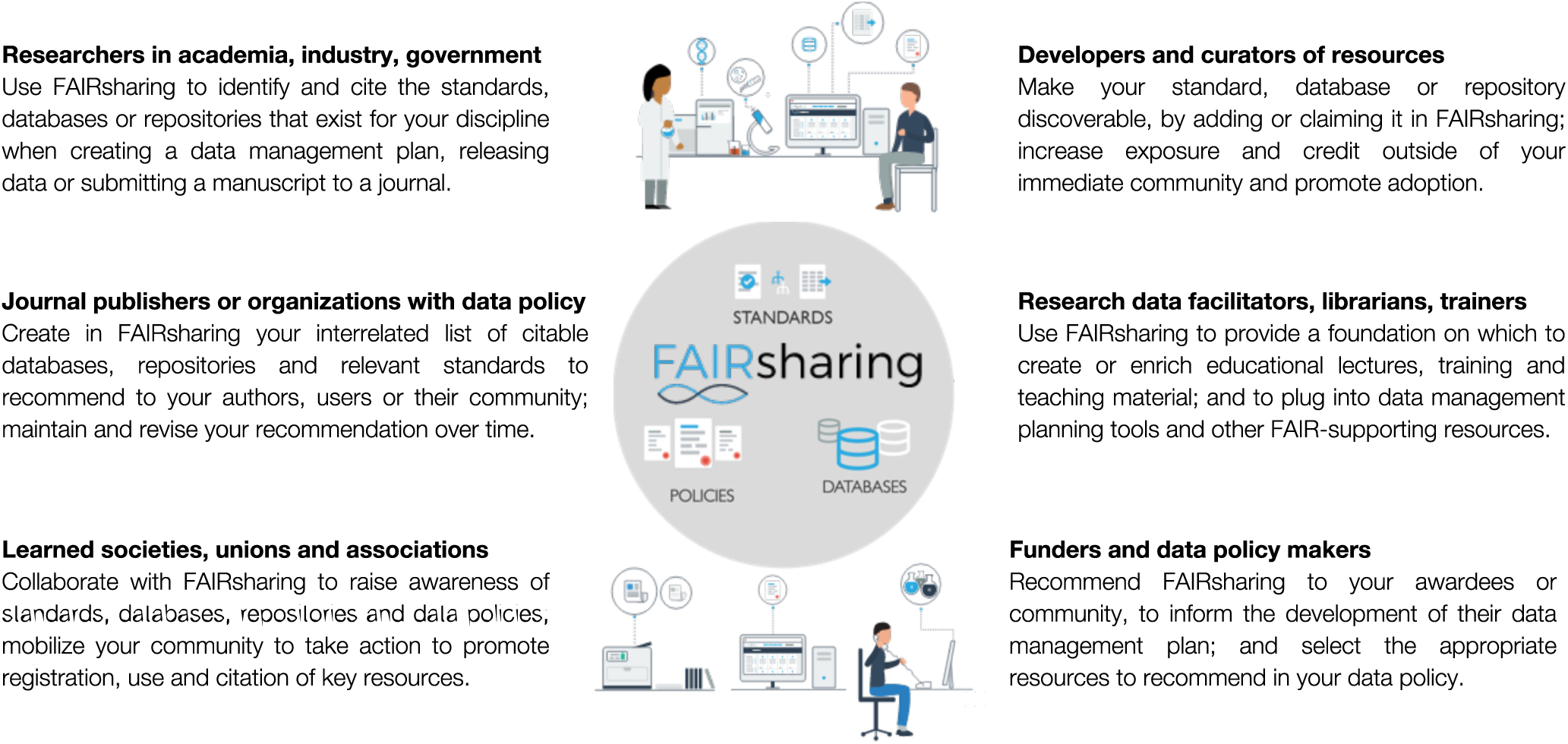
Several stakeholders play catalytic roles to foster a culture change within the research community into one where the use of standards, databases and repositories for FAIRer data is pervasive; this figure summarizes FAIRsharing guidance to each each stakeholder group.

Standard developers and database curators can use FAIRsharing to explore what resources exist in their areas of interest (and if those resources can be used or extended), as well as enhance the discoverability and exposure of their resource. This resource might then receive credit outside of their immediate community and ultimately promote adoption (to learn how to add your resource to FAIRsharing, or to claim it, see https://fairsharing.org/new). A representative of a community standardization initiative is best placed to describe the status of a standard(s) and to track its evolution. This can be done by creating an individual record (e.g., the DDI standard for social, behavioral, economic, and health data; https://doi.org/10.25504/FAIRsharing.1t5ws6) or by grouping several records together in a collection (e.g., the HUPO PSI standards for proteomics and interactomics data; https://fairsharing.org/collection/HUPOPSI). To achieve FAIR data, linked data models need to be provided that allow the publishing and connecting of structured data on the web. Similarly, representatives of a database or repository are uniquely placed to describe their resource, and to declare the standards implemented (e.g., the ICPSR archive of behavioral and social science research data that uses the DDI standard: https://doi.org/10.25504/FAIRsharing.y0df7m) or the Reactome knowledge base, https://doi.org/10.25504/FAIRsharing.tf6kj8), which uses several standards in the COMBINE collection for computational models in biology networks: https://fairsharing.org/collection/ComputationalModelingCOMBINE). The more adopted a resource is, the greater its visibility. For example, if your standard is implemented by a repository, these two records will be interlinked; thus, if someone is interested in that repository they will see that your standard is used by that resource. If your resource is recommended in a data policy from a journal, funder or other organization, it will be given a ‘recommended’ ribbon, which is present on the record itself and clearly visible when the resource appears in search results.

For journal publishers or organizations with a data policy, FAIRsharing enables the maintenance of an interrelated list of citable standards and databases, grouping those that the policy recommends to users or their community (e.g., see examples of recommendations created by eight main publishers and journals: https://fairsharing.org/recommendations. including some generalist and many domain-specific databases and repositories). As we continue to map the landscape, journals/publishers can also revise their selections over time, enabling the recommendation of additional resources with more confidence. All journals that do not have such data statements should develop them to ensure all data relating to an article or project are as FAIR as possible. Finally, journals should also encourage authors to cite the standards, database and repositories they use or develop via the ‘how to cite this record’ statement, found on each FAIRsharing record, which includes a DOI.

Trainers, educators as well as librarians and those organization and services involved in supporting research data can use FAIRsharing to provide a foundation on which to create or enrich educational lectures, training and teaching material, and to plug it into data management planning tools. These stakeholder communities play a pivotal role to prepare the new generation of scientists and deliver courses and tools that address the need to guide or empower researchers to organize data and to make it FAIR.

Learned societies, international scientific unions and associations, and alliances of these organisations should raise awareness around standards, databases. repositories and data policies, in particular on their availability, scope and value for FAIR and reproducible research; as well as mobilize their community members to take action^e.g.10^,^11^,^12^, to promote the use and adoption of key resources, initiate new or participate in existing initiatives to define and implement policies and projects.

Funders can use FAIRsharing to help select the appropriate resources to recommend in their data policy and highlight those resources that awardees should consider when writing their data management plan^e.g.13^. If we are to make FAIR data a reality, funders should recognize standards, as well as databases and repositories, as digital objects in their own right, which have and must have their own associated research, development and educational activities^14^. New funding frameworks need to be created to provide catalytic support for the technical and social activities around standards, in specific domains, within and across disciplines to enhance their implementation in databases and repositories, and the interoperability and reusability of data.

Last but not least, researchers can use FAIRsharing as a lookup resource to identify and cite the standards, databases or repositories that exist for their data and discipline, for example, when creating a data management plan for a grant proposal or funded project; or when submitting a manuscript to a journal, to identify the recommended databases and repositories, as well as the standards they implement to ensure all relevant information about the data is collected at the source. Today’s data-driven science, as well as the growing demand from governments, funders and publishers for FAIRer data, requires greater researcher responsibility. Acknowledging that the ecosystem of guidance and tools is still work in progress, it is essential that researchers develop or enhance their research data management skills, or seek the support of professionals in this area.

Anyone can be a user of FAIRsharing. FAIRsharing brings the producers and consumers of standards, databases, repositories and data policies closer together, with a growing list of adopters (https://fairsharing.org/communities). Representatives of institutions, libraries, journal publishers, funders, infrastructure programmes, societies and other organizations or projects (that in turn serve and guide individual researchers or other stakeholders on research data management matters) can become an adopter.

Help us to help you. We welcome collaborative proposals from complementary resources, we are open to participate in joint projects to develop services for specific stakeholders and communities. Join us or reach out to us, and let’s pave the way for FAIRer data together.

## Acknowledgements

We thank all our users, producers and consumers of standards, databases, repositories and policies, as well as adopters, past advisory board members, collaborators and content contributors to the current resource and its precursors (BioSharing and the Minimum Information about a Biomedical or Biological Investigation, MIBBI portal). We specifically thank key past contributors to the resource, notably Eamonn Maguire, Annapaola Santarsiero, and Chris Taylor. We are also very grateful to teams at the British Library and the University of Oxford’s Bodleian Libraries for their continued support with minting DOIs. Some of the discussion in this article and call for action was developed as part of the joint RDA and Force11 working group; therefore, we acknowledge the support provided by the RDA and the Force11 communities and structures. The main authors are funded by grants awarded to S.A.-S. that include elements of FAIRsharing; specifically grants from the UK BBSRC and Research Councils (BB/L024101/1, BB/L005069/1), EU (H2020-EU.3.1, 634107, H2020-EU.1.4.1.3, 654241, H2020-EU.1.4.1.1, 676559), IMI (116060), and NIH (U54 AI117925, 1U24AI117966-01, 1OT3OD025459-01, 1OT3OD025467-01, 1OT3OD025462-01) and the new FAIRsharing award from the Wellcome Trust (212930/Z/18/Z) as well as a related one (208381/A/17/Z). S.A.-S. is funded also by the Oxford e-Research Centre, Department of Engineering Science of the University of Oxford.

## Authors contributions

The core authors represent the initial operational team of the FAIRsharing resource. S-A.S. developed the concept and provided the strategic direction, and with P.R.-S. launched the initial portal in 2011, which was re-branded, curated, enriched and further developed by A.L., M.T., M.I. and A.G.-B. under the coordination of P.McQ. Under the FAIRsharing Community, we list the new members of the operational team, D.B. and R.G., key content curators, M.A. and D.D., as well as advisory board co-chairs, E.G. and V.K., special advisor, J.T., and Force11 and the RDA working group co-chairs, S.H. and R.L., followed by core adopters, advisory board members, and/or key collaborators and contributors to the collections and recommendations. P.McQ. assembled the statistics, and S-A.S. and P.McQ wrote the manuscript with contributions from the core authors and approval of the FAIRsharing Community.

